# NEUROMETABOLOMIC IMPACTS OF MODELED WILDFIRE SMOKE AND PROTECTIVE BENEFITS OF ANTI-AGING THERAPEUTICS IN AGED FEMALE C57BL/6J MICE

**DOI:** 10.1101/2023.09.21.558863

**Authors:** David Scieszka, Haiwei Gu, Amanda Barkley-Levenson, Ed Barr, Marcus Garcia, Jessica G Begay, Guy Herbert, Kiran Bhaskar, Mark McCormick, Jonathan Brigman, Andrew Ottens, Barry Bleske, Matthew J Campen

**Author notes:** To whom correspondence should be addressed. Matthew J Campen, PhD, Department of Pharmaceutical Sciences, MSC09 5360, 1 University of New Mexico, Albuquerque, NM 87131-0001. **Author Contributions:** Design of Experiments – MJC, DS, ABL, AO, BB, JB; Conduct of experiments – DS, HG, ABL, EB, MG, JGB, GH, MJC; Analysis of Data – DS, MJC; Writing – DS, MJC, KB, MM, JB, AO, BB.

## Abstract

Wildland fires have become progressively more extensive over the past 30 years in the US, and now routinely generate smoke that deteriorates air quality for most of the country. We explored the neurometabolomic impact that smoke derived from biomass has on older (18 months) female C57BL/6J mice, both acutely and after 10 weeks of recovery from exposures. Mice (N=6/group) were exposed to wood smoke (WS) 4 hours/day, every other day, for 2 weeks (7 exposures total) to an average concentration of 0.448mg/m^3^ per exposure. One group was euthanized 24 hours after the last exposure. Other groups were then placed on 1 of 4 treatment regimens for 10 weeks after wood smoke exposures: vehicle; resveratrol in chow plus nicotinamide mononucleotide in water (RNMN); senolytics via gavage (dasatanib+quercetin; DQ); or both RNMN with DQ (RNDQ). Among the findings, the aging from 18 months to 21 months was associated with the greatest metabolic shift, including changes in nicotinamide metabolism, with WS exposure effects that were relatively modest. WS caused a reduction in NAD+ within the prefrontal cortex immediately after exposure and a long-term reduction in serotonin that persisted for 10 weeks. The serotonin reductions were corroborated by forced swim tests, which revealed an increased immobility (reduction in motivation) immediately post-exposure and persisted for 10 weeks. RNMN had the most beneficial effects after WS exposure, while RNDQ caused markers of brain aging to be upregulated within WS-exposed mice. Findings highlight the persistent neurometabolomic and behavioral effects of woodsmoke exposure in an aged mouse model.

**Significance Statement:** Neurological impacts of wildfire smoke are largely underexplored but include neuroinflammation and metabolic changes. The present study highlights modulation of major metabolites in the prefrontal cortex and behavioral consequences in aged (18 month) female mice that persists 10 weeks after wood smoke exposure ended. Supplements derived from the anti-aging field were able to mitigate much of the woodsmoke effect, especially a combination of resveratrol and nicotinamide mononucleotide.

## INTRODUCTION

Exposure to wildfire smoke is becoming a growing concern for the global population, particularly as this population ages. Over the past 15 years, there has been a documented increase in the extent of wildfires and resulting damage in the United States, which is correlated with changing global temperatures(1). In 2020, the wildfires on the west coast generated smoke that spread high concentrations of particulate matter (PM) throughout much of the United States, potentially affecting the health of hundreds of millions of people for several months. Wildfires release toxic airborne particulates and gases from varying sources, including biomass and human-made fuels, which can cause harm to populations far from the fire source. While many combustion-derived gases remain localized, both PM and carbon monoxide are stable components of wildfire smoke that can travel long distances, which partially explains the increase in hospital admissions for respiratory issues in states outside the wildfire smoke source(2–5). Finally, it should be noted that there is a growing concern for minority populations, including Native American communities, which are at a disproportionately higher risk of wildfires than other communities(6–8).

The complex mixture of inhaled toxicants from wildfires can harm not only the lungs, but also cause systemic health effects(9). Yet, the long-term and short-term effects of wildfire smoke-derived PM on neurological outcomes, including its role in age-related diseases, are not well understood. Independent of wildfires, PM_2.5_ (particles less than 2.5 microns in diameter) has been linked to a range of adverse neurological outcomes, including Alzheimer’s disease related dementias (ADRD), suicide, depression, psychosis, and others(10–13). Impairment of the blood-brain barrier and resultant neuroinflammation are believed to play a role in these neurological disorders(14, 15). The blood-brain barrier is made up of tightly connected brain endothelial cells that line the central nervous system’s blood vessels, surrounded by astrocytic end feet, and monitored by nearby microglia. Recent research suggests that inhaled PM can cause proteolytic activity in the lung, leading to the release of bioactive peptide fragments into the bloodstream that specifically target endothelial cells and promote neuroinflammation(16), lead to infiltration of peripheral immune cells, and alter cytokine profiles(17, 18).

Wildfire smoke has been associated with pulmonary senescence fates(19) as well as adverse neurological outcomes arising from inflammation(17, 20). Pharmacological interventions hold promise for cognitive benefits and cardioprotective effects, which could offset wildfire smoke-induced neurological and cardiovascular conditions(21). Senescence is a proinflammatory state of cell cycle arrest that has been hypothesized as a root cause of aging(22–25), and can cause systemic inflammation through the release of senescence-associated secretory phenotype molecules into the circulation(26). More specifically, the senescence-associated secretory profile (SASP) is characterized by the release of proinflammatory molecules, including cytokines and interleukins, into the extracellular space(26, 27). To combat this release, senolytics are a class of naturally occurring drugs that target and block senescent cell anti-apoptotic pathways, allowing for apoptosis to occur(28). If allowed to persist, this SASP-associated inflammation has been linked with type 2 diabetes incidence(29), cardiovascular disease(30), cancer(31), accelerated brain aging(32), and neurodegenerative disorders(33), among others(34). Resveratrol is a natural polyphenol, abundant in red wine, that has shown effects as an anti-inflammatory, cancer management, neuroprotective, and cardioprotective molecule(35–37), with many of these effects arising from the activation of sirtuin 1 and other nicotinamide adenine dinucleotide (NAD^+^)-consuming enzymes. Nicotinamide mononucleotide (NMN) is a NAD^+^ precursor(38), whose supplementation has been shown as beneficial for cellular NAD^+^ abundance, longevity(39), cognition(40), ADRD(41), and depression(42), among others(43).

Many investigations have reported antioxidant effects from supplementation with resveratrol(44, 45) and nicotinamide mononucleotide (NMN)(46, 47, 47), and healthspan benefits from the combination of senolytics dasatinib + quercetin (DQ)(48–50). However, only recently have studies been performed on the combination of resveratrol with NMN (RNMN)(51), and no studies have combined resveratrol, NMN, dasatinib, and quercetin (RNDQ). The introduction of DQ could target WS-associated senescent cells, while RNMN could boost cardio- and neuroprotection. It is known that the benefits of DQ are additive(52). Therefore, we examined whether the combinations of RNMN, and the more complete cocktail of RNDQ would elicit additive benefits in terms of diminishing negative impacts of laboratory-generated woodsmoke (WS) inhalation on neurological health, and possibly outcomes simply related to advanced aging.

## RESULTS

### Experimental Design and Exposure Characterization

Average concentration per exposure was calculated to be 0.448 mg PM/m^3^ for each of the 7 x 4h exposures (Fig. 1B), with a 24-hour exposure concentration (in line with U.S. Environmental Protection Agency (EPA) methods for particulate matter (PM) regulations of 0.037 mg PM/m^3^. We measured the levels of carbon monoxide (CO) and oxides of nitrogen species (NOx) for all exposures. They were well below EPA National Ambient Air Quality Standard levels(Fig. 1C). The 8h average allowable limit for CO is 9 parts per million (PPM) and for a single hour of exposure the concentration limit is 35 PPM. Similarly, the 1-hour average allowable limit for NO_x_ is 100 parts per billion (PPB), and the 1-year average exposure limit is 53 PPB (53, 54). Of the 40 total 18-month-old mice exposed to WS, only one died during the 14d exposure period, while no FA controls died. In the subsequent 10 weeks of pharmacological treatment, 2 of the WS mice (out of 34) died before study completion, both in the RNDQ treatment arm; no FA mice died prematurely.

**Figure 1.**
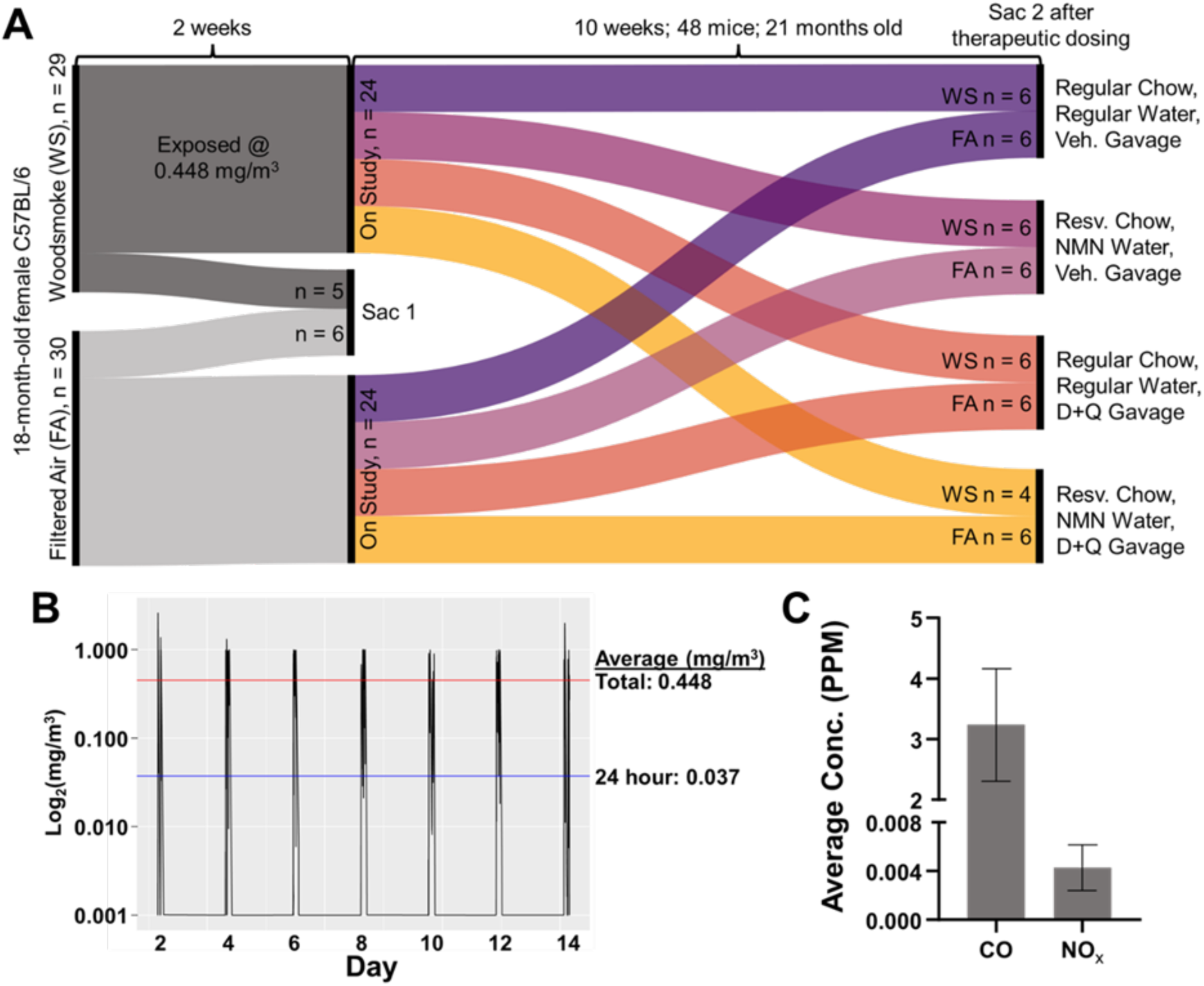
Schematic of experimental design and exposure characterization. A) Experimental design schematic. 18-month-old female C57BL/6J mice were exposed to WS at an average concentration of 0.448mg/m3 every other day for 2 weeks (7 total exposures). 1 day post exposures, one cohort of FA and WS were euthanized. The remaining mice went on study for 10 weeks in their individual drug groups. At the end of therapeutic dosing, each cohort was euthanized for metabolomic sequencing. B) Concentration measuring during each 4-hour exposure. Top line (red): average across all exposures. Bottom line (blue): average across all exposures, taking 24 hours per day into account. C) Levels of carbon monoxide (CO) and oxides of nitrogen (NOX). PPM: parts per million.

### Untargeted Metabolomics

#### Overall

The initial data analysis assessed broad swaths of differences between groups (Fig. 2A). There were significant metabolomic differences between 18-month-old FA and WS in the PFC, indicating an exposure effect. The natural course of aging was also calculated as significant, through the comparisons of 18mo Veh to 21mo Veh for both FA and WS, and through the grouped comparison of FA&WS 18mo to FA&WS 21mo. The RNMN intervention group trended differently between 21mo FA vs WS exposure conditions and was significantly different between exposure-matched WS 21mo Veh. FA RNMN vs FA Veh showed no differences. However, the FA DQ group was significantly different compared to age-matched FA Veh. These findings indicated that the combination of DQ had the strongest effect on the overall PFC metabolic profile of unexposed mice, while RNMN had the strongest effect on WS-exposed mice.

**Figure 2.**
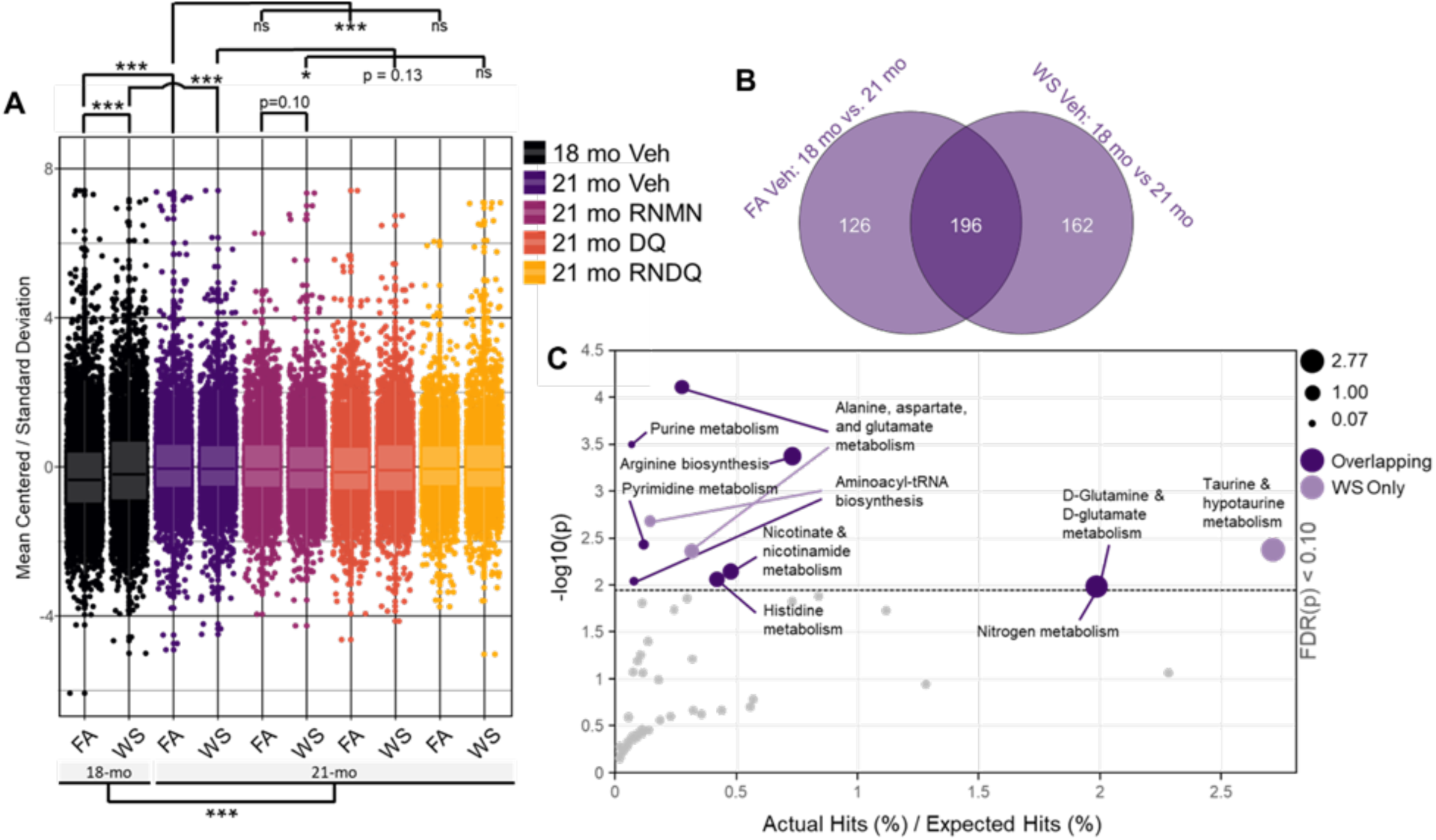
Initial difference testing and metabolomic aging verification. A) Linear mixed effects regression modeling the entire metabolomic dataset. 18mo mice showed immediate exposure-induced differences. 21mo mice showed differences between their 18mo counterparts for both conditions. B) Venn diagram of statistically significant metabolites. More metabolites were overlapping, indicating an aging effect was more prominent than the WS effects, but that WS exposure had a lasting outcome (student’s t-tests). C) Significant pathways affected by metabolites that overlapped (dark purple) and those that were exclusive to WS (light purple). The nicotinamide and tRNA perturbations were largely confirmatory of an aging phenotype. The alteration to neuroprotective taurine indicates a risk to oxidative damage. Combination of FA Veh and overlap resulted in no additional significant pathways.

#### Aging Influence on PFC Metabolites

To interrogate the natural aging effect further, significant metabolite differences from 18 months to 21 months, for both FA and WS groups, were compiled into lists and compared as a Venn diagram (Fig 2B). We observed a greater number of significant overlapping metabolites (196) than significant non-overlapping metabolites in FA (126) or WS (162). The overlapping cluster reflects aging-related changes that are unperturbed by WS exposure. We extracted the significant overlapping metabolites and input them into pathway analysis. After FDR correction (cut-off p<0.1), the main pathways affected included alanine, aspartate, and glutamate metabolism; purine biosynthesis; arginine biosynthesis; pyrimidine metabolism; and nicotinate and nicotinamide metabolism (Fig 2C, Table 1). The affected pathway of nicotinate and nicotinamide metabolism confirms the accuracy of pathway analysis employed, based on abundant literature showing that aging is associated with declines in NAD^+^ reserves and biosynthesis (55–65). The non-overlapping FA metabolites were added into the list containing overlapping metabolites and the new list was queried for pathway alterations. Due to the number of metabolites entered into the software, FDR correction returned no significantly altered pathways for these metabolites altered in FA controls from age 18-21 months. Finally, we took the non-overlapping WS metabolites and queried them for pathway alterations. Previous work from our lab showed that wildfire exposures decreased neuroprotective taurine (17), which was confirmed here as one of the main affected pathways: taurine and hypotaurine metabolism. Taken together, these results indicate that the effect of aging over this 10-week period in mice has a greater impact on the metabolic profile than the effects of our subchronic WS exposure. Additionally, aging had a greater impact than any drug combination. However, the modest WS exposure levels still affected the metabolic profile after 10 weeks of natural recovery; and the drug combinations most affecting PFC’s of FA and WS are DQ and RNMN, respectively.

**Table 1.** Pathways resulting from Figure 2B. Overlapping metabolites of shared Venn diagram region (196 metabolites) and WS-exclusive (162 metabolites).

| Overlap |  |  |  |  |  |  |  |
| --- | --- | --- | --- | --- | --- | --- | --- |
| Pathway | Total | Hits | Expected | Impact | Raw p | $-\log_{10}(p)$ | FDR |
| Alanine, aspartate and glutamate metabolism | 28 | 6 | 0.781 | 0.274 | 0.000 | 4.11 | 0.007 |
| Purine metabolism | 66 | 8 | 1.841 | 0.066 | 0.000 | 3.50 | 0.012 |
| Arginine biosynthesis | 14 | 4 | 0.390 | 0.732 | 0.000 | 3.37 | 0.012 |
| Pyrimidine metabolism | 39 | 5 | 1.088 | 0.118 | 0.004 | 2.42 | 0.079 |
| Nicotinate and nicotinamide metabolism | 15 | 3 | 0.418 | 0.478 | 0.007 | 2.14 | 0.099 |
| Histidine metabolism | 16 | 3 | 0.446 | 0.420 | 0.009 | 2.06 | 0.099 |
| Aminoacyl-tRNA biosynthesis | 48 | 5 | 1.339 | 0.078 | 0.009 | 2.03 | 0.099 |
| Nitrogen metabolism | 6 | 2 | 0.167 | 1.992 | 0.011 | 1.97 | 0.099 |
| D-Glutamine and D-glutamate metabolism | 6 | 2 | 0.167 | 1.992 | 0.011 | 1.97 | 0.099 |
| Pantothenate and CoA biosynthesis | 19 | 3 | 0.530 | 0.298 | 0.014 | 1.84 | 0.120 |
| Glycerophospholipid metabolism | 36 | 4 | 1.004 | 0.111 | 0.016 | 1.79 | 0.123 |
| beta-Alanine metabolism | 21 | 3 | 0.586 | 0.244 | 0.019 | 1.72 | 0.124 |
| Valine, leucine and isoleucine biosynthesis | 8 | 2 | 0.223 | 1.121 | 0.019 | 1.72 | 0.124 |
| Glutathione metabolism | 28 | 3 | 0.781 | 0.137 | 0.041 | 1.39 | 0.244 |
| Glyoxylate and dicarboxylate metabolism | 32 | 3 | 0.892 | 0.105 | 0.057 | 1.24 | 0.320 |
| Butanoate metabolism | 15 | 2 | 0.418 | 0.319 | 0.063 | 1.20 | 0.328 |
| Glycine, serine and threonine metabolism | 34 | 3 | 0.948 | 0.093 | 0.066 | 1.18 | 0.328 |
| Arginine and proline metabolism | 38 | 3 | 1.060 | 0.074 | 0.087 | 1.06 | 0.404 |
| Ether lipid metabolism | 20 | 2 | 0.558 | 0.179 | 0.105 | 0.98 | 0.465 |
| Taurine and hypotaurine metabolism | 8 | 1 | 0.223 | 0.560 | 0.203 | 0.69 | 0.852 |
| One carbon pool by folate | 9 | 1 | 0.251 | 0.443 | 0.225 | 0.65 | 0.860 |
| Vitamin B6 metabolism | 9 | 1 | 0.251 | 0.443 | 0.225 | 0.65 | 0.860 |
| Ascorbate and aldarate metabolism | 10 | 1 | 0.279 | 0.359 | 0.247 | 0.61 | 0.892 |
| Biosynthesis of unsaturated fatty acids | 36 | 2 | 1.004 | 0.055 | 0.266 | 0.58 | 0.892 |
| Arachidonic acid metabolism | 36 | 2 | 1.004 | 0.055 | 0.266 | 0.58 | 0.892 |
| Glycerolipid metabolism | 16 | 1 | 0.446 | 0.140 | 0.365 | 0.44 | 1.000 |
| Fructose and mannose metabolism | 18 | 1 | 0.502 | 0.111 | 0.401 | 0.40 | 1.000 |
| Citrate cycle (TCA cycle) | 20 | 1 | 0.558 | 0.090 | 0.434 | 0.36 | 1.000 |
| Pyruvate metabolism | 22 | 1 | 0.614 | 0.074 | 0.466 | 0.33 | 1.000 |
| Glycolysis / Gluconeogenesis | 26 | 1 | 0.725 | 0.053 | 0.524 | 0.28 | 1.000 |
| Folate biosynthesis | 27 | 1 | 0.753 | 0.049 | 0.537 | 0.27 | 1.000 |
| Galactose metabolism | 27 | 1 | 0.753 | 0.049 | 0.537 | 0.27 | 1.000 |
| Metabolism of xenobiotics by cytochrome P450 | 64 | 2 | 1.785 | 0.018 | 0.541 | 0.27 | 1.000 |
| Inositol phosphate metabolism | 30 | 1 | 0.837 | 0.040 | 0.576 | 0.24 | 1.000 |
| Porphyrin and chlorophyll metabolism | 30 | 1 | 0.837 | 0.040 | 0.576 | 0.24 | 1.000 |
| Cysteine and methionine metabolism | 33 | 1 | 0.920 | 0.033 | 0.611 | 0.21 | 1.000 |
| Amino sugar and nucleotide sugar metabolism | 37 | 1 | 1.032 | 0.026 | 0.653 | 0.18 | 1.000 |
| Valine, leucine and isoleucine degradation | 40 | 1 | 1.116 | 0.022 | 0.682 | 0.17 | 1.000 |
| Tyrosine metabolism | 42 | 1 | 1.171 | 0.020 | 0.700 | 0.15 | 1.000 |
| Primary bile acid biosynthesis | 46 | 1 | 1.283 | 0.017 | 0.733 | 0.13 | 1.000 |

WS Only
| Pathway | Total | Hits | Expected | Impact | Raw p | $-\log_{10}(p)$ | FDR |
| --- | --- | --- | --- | --- | --- | --- | --- |
| Aminoacyl-tRNA biosynthesis | 48 | 4 | 0.542 | 0.154 | 0.002 | 2.80 | 0.092 |
| Taurine and hypotaurine metabolism | 8 | 2 | 0.090 | 2.768 | 0.003 | 2.49 | 0.092 |
| Alanine, aspartate and glutamate metabolism | 28 | 3 | 0.316 | 0.339 | 0.003 | 2.48 | 0.092 |
| Arginine biosynthesis | 14 | 2 | 0.158 | 0.904 | 0.010 | 2.00 | 0.194 |
| Starch and sucrose metabolism | 15 | 2 | 0.169 | 0.787 | 0.012 | 1.94 | 0.194 |
| D-Glutamine and D-glutamate metabolism | 6 | 1 | 0.068 | 2.461 | 0.066 | 1.18 | 0.698 |
| Nitrogen metabolism | 6 | 1 | 0.068 | 2.461 | 0.066 | 1.18 | 0.698 |
| Arginine and proline metabolism | 38 | 2 | 0.429 | 0.123 | 0.066 | 1.18 | 0.698 |
| Valine, leucine and isoleucine biosynthesis | 8 | 1 | 0.090 | 1.384 | 0.087 | 1.06 | 0.812 |
| Phenylalanine metabolism | 12 | 1 | 0.135 | 0.615 | 0.128 | 0.89 | 1.000 |
| Glycerolipid metabolism | 16 | 1 | 0.181 | 0.346 | 0.167 | 0.78 | 1.000 |
| Pantothenate and CoA biosynthesis | 19 | 1 | 0.214 | 0.245 | 0.195 | 0.71 | 1.000 |
| Sphingolipid metabolism | 21 | 1 | 0.237 | 0.201 | 0.213 | 0.67 | 1.000 |
| Galactose metabolism | 27 | 1 | 0.305 | 0.122 | 0.266 | 0.58 | 1.000 |
| Glutathione metabolism | 28 | 1 | 0.316 | 0.113 | 0.274 | 0.56 | 1.000 |
| Glyoxylate and dicarboxylate metabolism | 32 | 1 | 0.361 | 0.087 | 0.307 | 0.51 | 1.000 |
| Cysteine and methionine metabolism | 33 | 1 | 0.373 | 0.081 | 0.315 | 0.50 | 1.000 |
| Glycine, serine and threonine metabolism | 34 | 1 | 0.384 | 0.077 | 0.323 | 0.49 | 1.000 |
| Glycerophospholipid metabolism | 36 | 1 | 0.406 | 0.068 | 0.339 | 0.47 | 1.000 |
| Pyrimidine metabolism | 39 | 1 | 0.440 | 0.058 | 0.361 | 0.44 | 1.000 |
| Valine, leucine and isoleucine degradation | 40 | 1 | 0.452 | 0.055 | 0.369 | 0.43 | 1.000 |
| Primary bile acid biosynthesis | 46 | 1 | 0.519 | 0.042 | 0.412 | 0.39 | 1.000 |
| Purine metabolism | 66 | 1 | 0.745 | 0.020 | 0.535 | 0.27 | 1.000 |

#### WS and Drug Interventions

We then sought to determine the WS and drug intervention impact on metabolic profile and asked which drug combination would have the weakest overlap with the comparison between FA vs WS: 21 mo Veh (Fig 3). The comparison of FA vs WS at 21mo timepoint should reveal the metabolic profile skewed by WS, rather than the natural effects of aging. A weak overlap between a drug combination with the WS profile could indicate a metabolic shift away from the deleterious exposure outcomes. The comparisons examined FA 21mo vs WS 21mo mice, with significant metabolites totaling 270 for Veh, 22 for RNMN, 47 for DQ, and 49 for RNDQ. Venn diagrams revealed the weakest overlap between Veh and RNMN (3 exclusive + 1 shared with DQ), middle overlap of Veh with RNDQ [13], and strongest overlap between Veh and DQ [29] (Fig 3A). However, these overlaps were relatively small compared to the overall outcomes of FA vs WS Veh at the 21mo timepoint [270]. Relative to the other conditions, this indicated that DQ was least able to shift metabolic profile away from the WS exposure profile, with the combination of RNMN having the greatest shift after WS exposure. We also employed a Jaccard calculation to remove weighted bias based on the total number of significantly altered metabolites per condition. Regardless of methodology employed, we see the strongest overlap with Veh and DQ (J_index_: 0.08517), a middle overlap of Veh with RNDQ (0.0376), and the weakest overlap of Veh with RNMN (0.0103) (Fig. 3B). These indicated a minor reduction in effectiveness of drug combinations DQ and RNDQ after WS exposure and prompted a thorough investigation into individual metabolite contributions.

**Figure 3.**
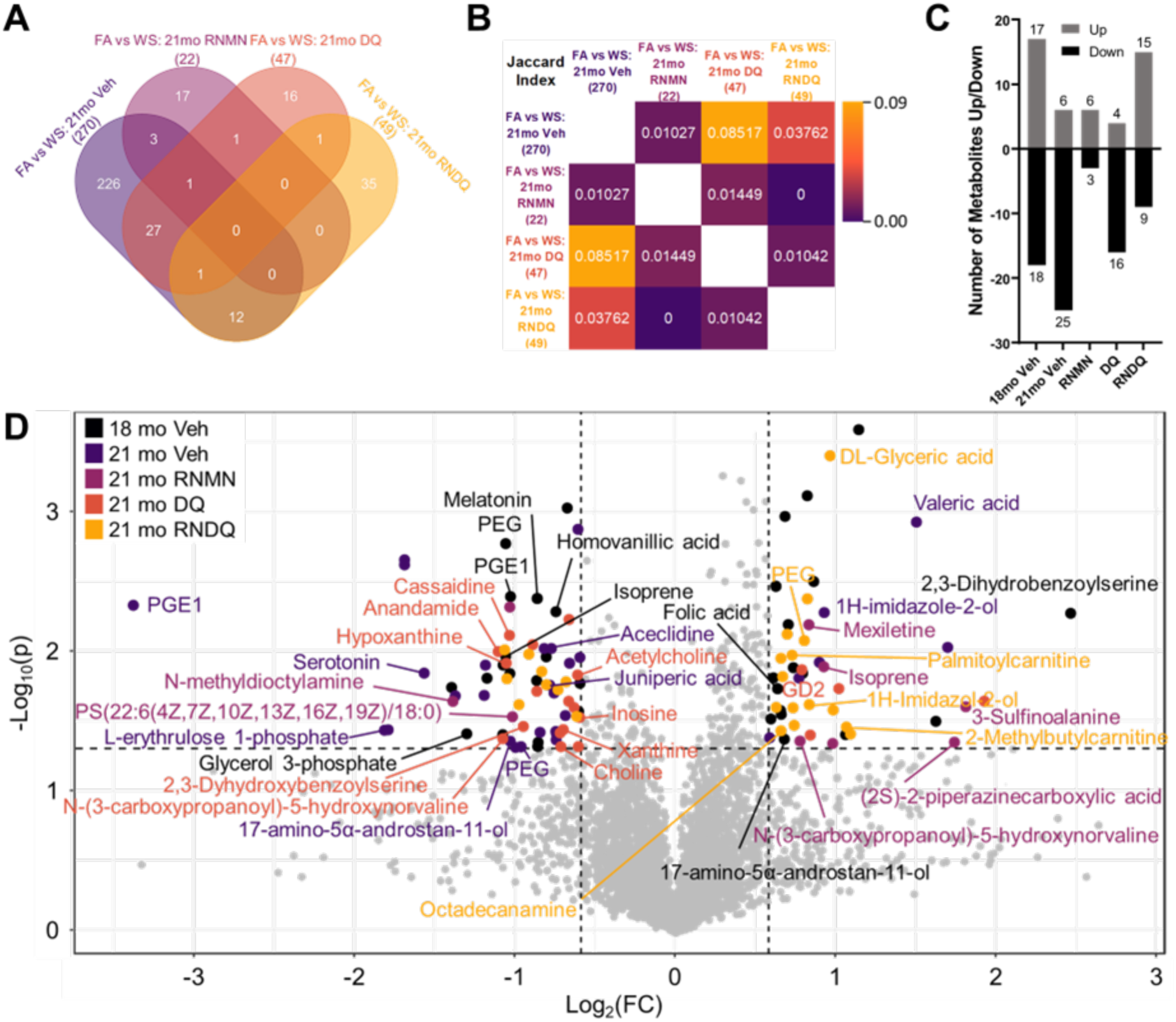
Drug correlations and overlay of significantly altered metabolites. A) Venn diagram of statistically significant metabolites. The metabolic differences between FA and WS Veh at 21mo illustrate a skewed metabolic profile the was unable to be resolved after 10 weeks (dark purple). These are compared to significantly different metabolites for each drug combination. The condition most closely related to no intervention (Veh) was DQ (78 overlapping), followed by RNDQ (18), and RNMN (6; student’s t-tests). B) For each group, Jaccard indices were calculated to reduce number weight bias. Still, DQ was most similar to Veh (no intervention), followed by RNDQ and RNMN, but RNMN and RNDQ were much closer to each other. C) Significant volcano-plot-calculated metabolites up/down per condition comparing FA versus WS. Of all the profiles, RNDQ appeared most similar to 18mo Veh. D) Overlay of untargeted and NAD+ metabolomic panels. Black: young, 18mo-old mice. Dark purple: natural course of aging for 10 weeks after WS exposure without drug intervention. Light purple: resveratrol and NMN. Orange: dasatinib and quercetin (DQ). Yellow: RNMN + DQ (RNDQ).

Volcano plots were compiled for drug-age-matched exposure condition (FA vs WS). The number of significantly up and downregulated metabolites revealed several patterns (Fig. 3C). Both ages of (FA vs WS) Veh had the largest number of metabolites that were significantly *decreased* (18mo: 18; 21mo: 25), while 18mo Veh and RNDQ had the greatest number that were significantly *increased* (18mo: 17; RNDQ:15; Fig. 3C). Of the drugs, RNDQ had the largest total of significantly altered metabolites in the FA vs WS condition [24] while RNMN had the least number of significantly altered [9]. Together, these indicated an inability to fully resolve WS exposure over 10 weeks without intervention, and that RNMN most effectively resolved WS changes while RNDQ elicited the greatest differential drug response between FA and WS. Taken together, these data illustrate the differences seen at each level of examination (with and without corrections). Without FDR correction, Venn diagrams revealed broad metabolic profiles that explained the aging effect (Figs. 2B, 3A, B). With FDR correction, pathway analyses, lmer, and volcano plots revealed the metabolites most responsible for metabolic shifts (Figs. 2A, C; 3D).

Overall, the volcano plot for 18mo Veh illustrated a WS effect that is larger than expected from such a modest exposure (Fig. 3D). After 10 weeks, the 21mo Veh mice still had not fully resolved this response to baseline. Of the drug combinations, RNDQ caused the upregulation of two metabolites known to exist within the aging murine brain (DL-glyceric acid(66) and octadecylamine(67)). This indicates a potential hazardous outcome for taking this drug regimen after WS exposure. Finally, RNMN appeared to recover the PFC to a greater extent than no intervention (21mo Veh), DQ on its own, and the combination of RNDQ (Fig. 3 C, D).

### Targeted Metabolomics

We examined the serotonin and dopamine pathways, as well as N-acetylaspartylglutamic acid (N-acetylaspartylglutamate or NAAG), GABA and NAD^+^. In the dopamine pathway, our untargeted panel was able to detect phenylalanine, tyrosine, tyramine, and homovanillic acid (Fig. 4A i-iv), but dopamine itself was unresolved. At 1-day post-exposure, 18mo Veh mice saw an upregulation to tyramine (Fig. 4A iii) and downregulation of homovanillic acid (Fig. 4A iv), which are the upstream and downstream metabolites of dopamine, respectively. The tripartite glutamine, glutamate, pyroglutamate pathway (Fig. 4B i-iii) illustrates an increase to glutamate at the 21mo time point. NAAG facilitates the release of glutamate (Fig. 4B iv) and was examined as a potential cause for upregulation of glutamate in 21mo Veh. From these 4 panels, it appears that 18mo Veh glutamate downward trends cannot be explained by other metabolites in this tripartite pathway. However, these data suggest that NAAG is partially responsible for the increases to glutamate and pyroglutamic acid in 21mo Veh. GABA was upregulated 1-day post exposure in the 18mo Veh animals, and we observed trending increases 10 weeks later in the 21mo Veh animals (Fig 4. C). NAD^+^ was decreased immediately after WS exposure, which indicates the potential for accelerated neurological aging (Fig. 4. D). Taken together, these data reveal long-term effects of WS on the levels of serotonin, glutamate, NAAG, and GABA (10 weeks), many of which are able to be resolved through the combinations of RNMN, DQ, or RNDQ.

**Figure 4.**
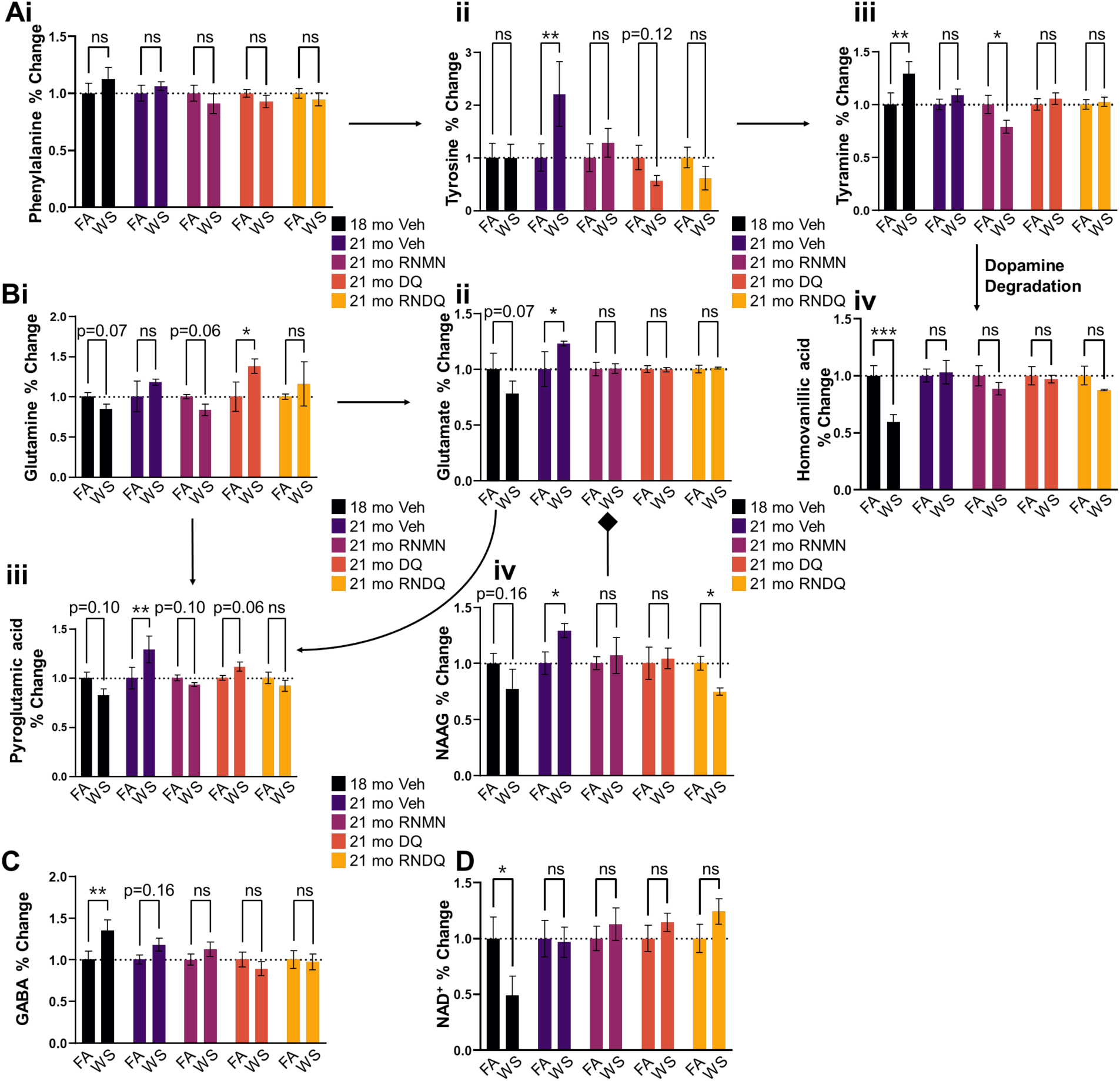
*De novo* neuro-metabolite investigations. Ai-iv) Metabolites detected within the dopamine synthesis and degradation pathway. iii) Downstream tyramine increased immediately after exposure. iv) Dopamine degradation frequency appears to be reduced based on lowered abundance of homovanillic acid. B i-iv) Metabolites detected in the tripart glutamine, glutamate, and pyroglutamic pathway along with NAAG, which facilitates the release of glutamate. Compensation mechanism could be in effect 10 weeks post exposure based on increased glutamate, NAAG, and pyroglutamic acid. C) GABA shows an immediate increase after exposure. D) NAD+ is decreased immediately post exposure.

We observed significant reductions in serotonin precursor, tryptophan (Fig. 5A i) at the 18mo time point, which was trending 10 weeks later in the 21mo Veh. These effects were mirrored in serotonin itself, with a trending reduction immediately post-exposure in the 18mo Veh and a significant decrease 10 weeks later in 21mo Veh animals (Fig. 5A ii). These data indicate a long-term downregulation of serotonin that is unable to be recovered naturally 10 weeks post exposure. To further examine the functional outcomes of observed serotonin reductions, we performed forced swim tests 1 day before exposures, 1 day after the last exposure, and after the 10-week recovery period (Figure 5B). When examining the pre-exposure timepoint, WS-exposed mice showed no significant differences in mobility when compared to FA mice. However, in all post-exposure cases — immediately after exposure, +5 weeks, and +10 weeks — immobility was observed as significantly higher in the WS group compared to FA controls, paralleling the sustained decreases in serotonin. Finally, we observed no difference in grip strength, illustrating that these observed effects were not the result of sarcopenia (Fig. 5C). Taken together, these data provide phenotypic evidence that the serotonin changes translated to depression-like behavior that persisted long after completion of exposures.

**Figure 5.**
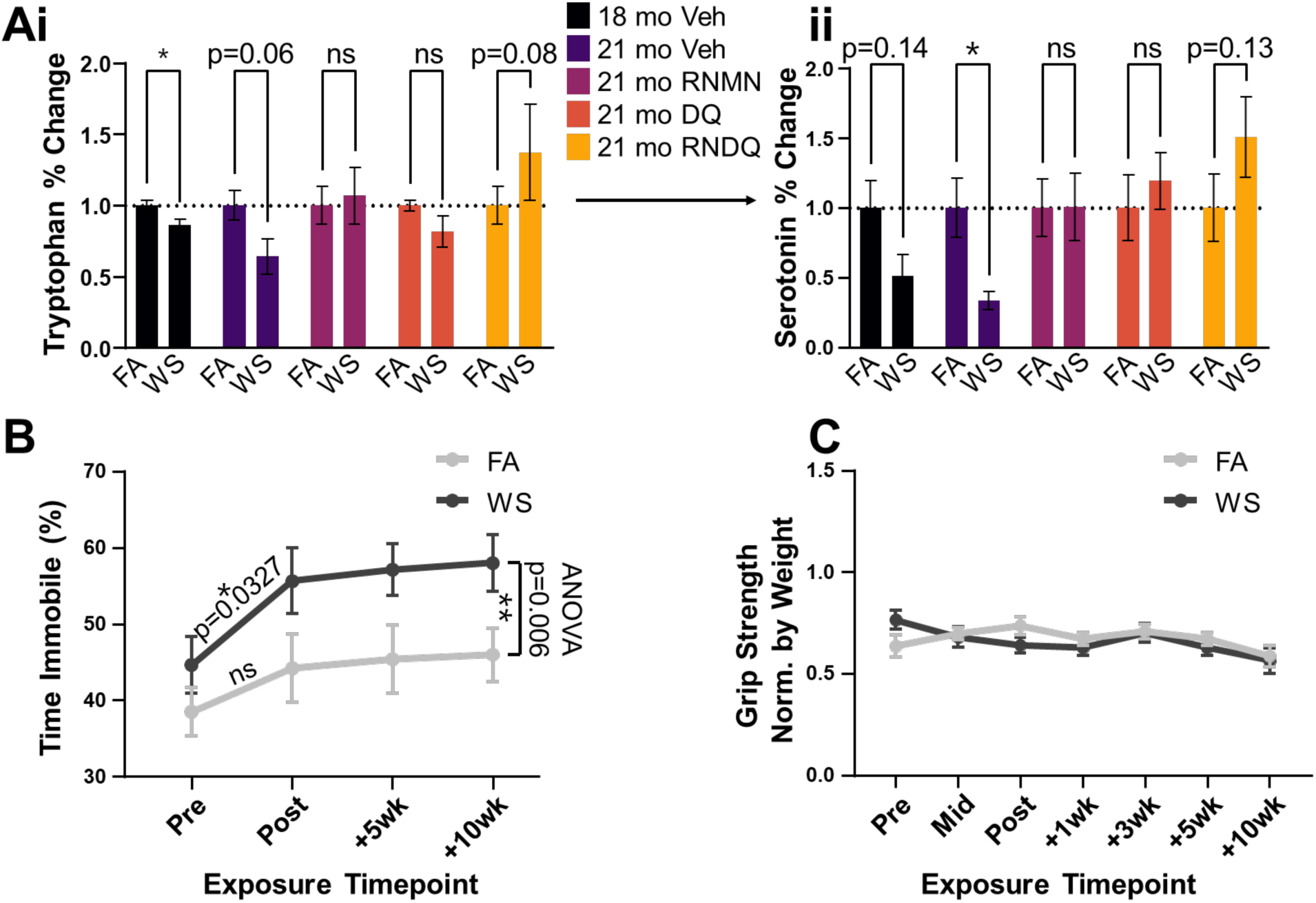
WS causes depression-like phenotype in mice. A i & ii) Metabolites detected in the serotonin synthesis pathway. i) Upstream tryptophan decreased immediately after exposure and trended decreased after 10 weeks of natural resolution. ii) Serotonin abundance trended decreased immediately after exposure but showed significantly decreased abundance after 10 weeks of natural resolution. B) Forced swim test corroborating serotonin reduction result, illustrating increased depression-like behavior. Initial test pre- vs post-exposure: student’s t-test. ANOVA test comparison for FA vs WS used all timepoints. C) Grip strength is not significantly different between groups and illustrates sarcopenia is not responsible for phenotypic outcomes observed.

## DISCUSSION

Broadly, this study examined the neuro-metabolomic effects after WS exposure in a model of advanced aging. Through the use of pharmacological interventions, we were able to query whether the current longevity drugs were more effective than natural resolution. While the neuro-metabolomic results broadly concur with recent studies in younger, 3-month-old animal models(18), the results could be seen as a poor predictor for the outcomes in aging models. Specifically, the persistence and magnitude of effects in the present study were notably extended relative to younger, 3-month-old animals(18). Critically, we observed a significant reduction to PFC serotonin that was unable to be naturally resolved in 10 weeks, and that resveratrol + NMN treatment was most beneficial after WS exposure compared to exposure-matched controls.

We examined WS because the total acres burned per year have roughly doubled over the past two decades (68). The concentrations of WS used are similar to earlier studies of naturally-occurring wildfire PM (17, 18), and well within range of human exposures that occur routinely during summer months (69). Expanding upon the most striking findings, WS exposure caused a sustained reduction in serotonin levels in the prefrontal cortex that persisted even after 10 weeks of recovery. The phenotypic expression of this change was supported by increased immobility in the forced swim test, suggesting the serotonin changes may have translated to increased depression-like behavior. These results parallel human studies showing a decreased ability to focus and learn after wildfire smoke exposure(70). Numerous epidemiological studies support a link between PM and neurological and behavioral health (10, 71, 72), but few have addressed wildfire smoke, specifically. A recent study, however, did note that risk of dementia from wildfire-derived PM was positive along with dusts from agricultural sources, while most other sources of PM were not associated with dementia outcomes (73).

Although many of the toxic gaseous components dissipate within close proximity to the fire source, the stability of carbon monoxide and PM_2.5_ mean that they can be transported great distances in the air. These components potentially cause adverse cardiovascular, pulmonary, and neurological outcomes thousands of kilometers away from the fire origins(17). Regardless of the wildfire point of origin, deleterious long-term outcomes have been implicated in exposure. These include neurological outcomes of depression, anxiety, suicide, ADRD, and decreased learning ability, as well as cardiac effects of stroke, heart disease, myocardial infarction, and atrial fibrillation, among other systemic effects as well(10–13, 15, 16, 27, 70).

Based on our experimental design (Fig 1A), we were able to query neurological effects of wildfire smoke and natural aging through metabolomic mass spectrometry. Previous work from our lab(17) and others(70, 74, 75) revealed metabolite alterations that were suggestive of mood alterations and impaired memory formation following WS exposure. Thus, we dissected the PFC to determine these effects and modeled them against aging, natural resolution of response, and the comparison of each drug regimen. To further assist in determining age-related effects, we employed a metabolomic panel designed to investigate the NAD^+^ synthesis pathway, and an additional untargeted panel. This experimental design was specifically chosen because it afforded the ability to examine a multitude of effects. Namely, the immediate exposure effects were determined through a comparison of FA vs WS at sac 1 (1-day post final exposure). We could determine natural resolution of inflammation over 10 weeks at sac 2 (after therapeutic dosing; dark purple). Through the combinations of different drug regimens, we could compare and contrast a multitude of effects, like natural aging, WS-associated aging, and drug rescuing effects, among others. For example, the difference between natural aging can be examined by comparing FA 18-month-old (18 mo) vehicle (Veh) to FA 21-month-old (21 mo) Veh. These naturally-altered age-associated metabolites can be compared to those found from WS 18 mo Veh vs Ws 21 mo Veh for overlap. The exact comparisons and their explanations are described below.

### FA vs WS: 18mo Veh – Immediate WS Exposure Effect

After 14 days of WS exposure, we saw changes to glycerol-3-phosphate (G3P), prostaglandin E1 (PGE1), homovanillic acid, and isoprene. G3P is used in the rapid replenishment of NAD^+^ stores in the brain and is reduced in our model. Reductions to PGE1 could indicate a request for platelet aggregation(76). Homovanillic acid is a major terminal metabolite of dopamine, with downregulation being associated with neuroinflammation and depression(77). Reduction to isoprene could indicate overuse of its antioxidant properties. This downregulation could also indicate oxidative damage undergoing repair. Together, the metabolites potentially reveal a landscape of inflammatory response and neurotransmitter modulation at 1-day post exposure with attempts to resolve inflammation.

### FA vs WS: 21mo Veh – Natural Resolution of Neural Metabolomic Changes

10 weeks following the WS exposure, prefrontal cortex PGE1 remained lower than in FA control mice, indicating a long-term exposure effect. Decreases to aceclidine could indicate dysregulation of cholinergic and dopaminergic neurotransmission, resulting in alterations to fear-extinguishing and working memory formation(78–80). Juniperic acid (16-hydroxyhexadecanoic acid) is an anti-inflammatory molecule, whose downregulation could indicate ongoing inflammatory responses that have not yet been resolved. Upregulation of valeric acid is known to worsen neuroinflammation, learning, and memory(81). Overall, we see metabolites that may indicate stress and decreases to memory formation after 10 weeks of attempted recovery from a modest WS exposure.

### FA vs WS: 21mo RNMN - Modulation of Resolution by RNMN

Mexiletine is upregulated and is known to protect against oxidative stress(82). 3-sulfoalanine upregulation could indicate NMDA receptor activation and an attempted recovery of dopaminergic-associated decision making and reward behavior. Although isoprene was downregulated at 18 months, we see it upregulated after the addition of RNMN, whose antioxidant effects could be helpful for the aging brain. 3-Sulfinoalanine is an NMDA agonist and catalyzes the irreversible transformation of kynurenine to kynurenic acid in activated astrocytes(83), which could stabilize neurons during times of stress. N-methyldiocylamine is similar to haloperidol(84), and improves thinking, mood, and behavior. Unfortunately for our mice, it was downregulated in both conditions. Notably, the downregulated phospholipid serine species (PS(22:6(4Z,7Z,10Z,13Z,16Z,19Z)/18:0)) is concentrated in myelin within brain tissue(85). 2-Piperazinecarboxylic acid has been used as an experimental treatment for anemia(86). Together, we see a mix of metabolites that have antioxidant and positive effects, with a sparse few that could negatively impact mood. Of all drug combinations, RNMN has the only metabolic profile indicating an overall positive outlook for brain health after WS exposure. However, many other changes from WS were resolved by different combinations of drugs, which supports the potential for a beneficial drug effect.

### FA vs WS: 21mo DQ – Modulation of Resolution by Senolytics

Upregulation of disialoganglioside 2 (GD2) has been shown to have anti-inflammatory effects in microglia (87). In conjunction with GD2, dihydroxybenzoylserine could have an anti-inflammatory effect. Downregulation of acetylcholine could be in response to vasodilation, or it could result in alterations to communication between the PFC and hippocampus (88). If communication is altered, it would reduce arousal, attention, and the ability to reinforce learning. Additionally, a significant reduction in choline was observed, which is a precursor to acetylcholine. Anandamide was downregulated, and plays a role in memory, sleep, appetite, and pain reductions. Hypoxanthine was reduced and is transported into the brain during times of purine shortages (89), indicating purine biosynthesis could be altered. Broadly, this aligns with pathway analysis that revealed alterations to purine metabolism during the natural course of aging. Hypoxanthine is a conversion product of xanthine, which was also reduced. Inosine was also reduced, which is a downstream product of hypoxanthine with a ribose ring. Overall, decreases to inosine, xanthine, and hypoxanthine are strong indicators of increased purine requirements within the PFC, which could be associated with a plethora of cellular processes. Additionally, metabolites indicate a reduction in arousal, attention, and decreased communication between the PFC and hippocampus.

### FA vs WS: 21mo RNDQ – Modulation of Resolution by Full Cocktail

After WS and RNDQ treatment, both DL-glyceric acid (66) and octadecylamine (67) were increased, and are both found in the aging mouse brain. Palmitoyl carnitine and 2-methylbutyrylcarnitine were both increased, indicating an increased need for ATP production. Overall, the RNDQ condition contains the only upregulated metabolites known to exist within the aging brain. Their existence implies a lack of effect for the combination of RNDQ that was not observed in the RNMN or DQ groups after WS exposure. From these, it appears that the differential effects of RNDQ could be beneficial in FA groups, but not after WS. This is further solidified by palmitoyl carnitine and 2-methylbutyrylcarnitine showing an increased energetic demand. It was notable that the full RNDQ was the only post-exposure treatment arm that experienced premature mortality. We could not conclude that this was due to drug effects or by chance, but clearly does not favor the use of this cocktail.

### Limitations and Final Thoughts

Several limitations for the study should be noted. As resveratrol activates NAD^+^ consuming enzymes for longevity-associated effects and NMN is a precursor to the enzymatic fuel source, NAD^+^, this combination was viewed strategically for the aging mouse model, but we did not assess the specific impact of either resveratrol or NMN in isolation. Additionally, the senolytic drugs dasatinib and quercetin have additive effectiveness, and are now administered in combination with fisetin for the most recent clinical trial. The combination of resveratrol, NMN, dasatinib, and quercetin appeared as a novel potential therapy, given their complementary individual effects on the aging pathways. Lastly, the choice of biomass (pinon wood) for this study was based on local fuel sources and may differ marginally from the numerous other sources. Importantly, wildfires are uncontrolled mixtures of grasses, shrubs, and trees, burning at different temperatures, with meteorological factors influencing transport, mixing, and atmospheric aging of PM. Thus, there may not be a perfect model for wildfire smoke exposure, given the variations of time, distance, and concentration to which millions of people are exposed. Finally, we specifically examined the PFC and made no comparisons to other regions of the brain. The pattern of metabolite alterations could be significantly different elsewhere in the brain, and warrants further investigation.

Summarily, these data reveal persistent alterations in the neurometabolome of aging mice following exposure to WS, with corroborating behavioral impacts. The WS exposure tended to promote or exacerbate the metabolomic changes associated with aging, suggesting that not only is aging a risk factor that promotes vulnerability to environmental contaminants, but that environmental contaminants may drive certain processes of aging. Findings help explain the epidemiologically-observed associations between PM exposure and neurological outcomes, and further highlight the need to better understand the neurological and psychological impacts that wildfire smoke may have on public health.

## MATERIALS AND METHODS

### Animals and WS Exposures

Female C57BL/6J mice (Jackson Labs) at 18 months of age were housed in AAALAC-approved facilities, on a 12h light:dark (7AM:7PM) cycle and allowed to acclimate for 1 week prior to experiments. Depending on experimental condition, they were provided either standard chow diet or resveratrol chow and DI water or nicotinamide mononucleotide (NMN) water (described below) ad libitum (Fig. 1A). A total of 60 mice were used, randomly and evenly divided into filtered air (FA) control and WS (WS) exposed groups; with 6 mice per reusable plastic animal case system (Fig. 1A). All procedures were conducted humanely with approval by the University of New Mexico Institutional Animal Care and Use Committee. For exposures, a BioSpherix Medium A-Chamber was used, with mice in reusable shoebox plastic animal case systems. They were outfitted with standard wire tops, and water was available to mice throughout the exposures, but food was withheld (4h/d).

Whole-body exposures to biomass combustion were conducted for 4 h every other day for 14 days (7 total exposures). Biomass smoke production was facilitated by a ceramic furnace encircling a quartz tube using chipped pinon wood as the fuel (18). This tube was connected to a dilution chamber, with subsequent plumbing into the exposure chamber (Supplemental Figure 1). Exposure chamber smoke abundance was facilitated by vacuum and/or pressurization, with total pressure monitoring to ensure min/max exposure chamber pressure never exceeded −/+ 25mm Hg. Concentrations were monitored in real-time and manually adjusted to provide consistent average exposure levels.

Exposure concentrations were measured with a with a DustTrak II (TSI, Inc; Shoreview, Minnesota) in real-time using a tube fed directly into the exposure chamber. 47 mm quartz filters were collected for the duration of each exposure, and gravimetrically confirmed final daily averages using a microbalance (XPR6UD5, Mettler Toledo) in a temperature-controlled laboratory. Particle size distribution was quantified using the TSI Laser Aerosol Spectrometer 3340A using a tube fed directly to the exposure chamber. Size distribution was acquired without mice in the exposure chamber, and distribution was measured over a single 2-hour run. Particle size distribution largely fell within PM _0-1_ (median range = 0.138-0.145 μm) with less than 1% of particles above PM_2.5_.

### Post-exposure Recovery and Drug Treatments

After the last round of exposures, one group of mice were euthanized (6 FA and 5 WS-exposed mice) with the remaining 24 mice per exposure group then randomized to 1 of the 4 drug regimens (Fig. 1A): (1) standard chow, deionized (DI) water, vehicle (Veh) gavage; (2) resveratrol milled into standard chow, NMN water, Veh gavage (RNMN); (3) standard chow, DI water, dasatinib and quercetin (DQ) gavage; (4) resveratrol chow, NMN water, DQ gavage (RNDQ). An additional group of mice was included to recapitulate metabolomic findings of group 1 (Veh control) in addition to applying a forced swim test and grip strength test to characterize potential behavioral implications. Resveratrol chow was achieved by milling 0.1% resveratrol by weight into standard chow (Envigo, WI, USA). Estimated intake of resveratrol was calculated as 5mg/30g mouse/day, or 167 mg/kg/day (90). NMN was added to water 1.3mg/mL per week. Estimated intake of NMN was calculated as 300mg/kg/day (91). Mice were gavaged with the formulation of Dasatinib (5 mg/kg) and Quercetin (10 mg/kg) in vehicle (60% Phosal, 10% ethanol, 30% PEG-400) or gavaged with vehicle alone. Phosal formulation was 50% phosphatidyl choline and 50% propylene glycol. Dasatinib and quercetin gavage stock was created the day of each administration. Drugs or vehicle were administered at 9-11am for 3 consecutive days per week (i.e., Monday-Wednesday), every other week, for the total period of 10 weeks (92).

### Forced Swim and Grip Strength Testing

A separate subset of mice underwent exposures to FA (N=10) or WS (N=10) as above and then administered a forced swim test and grip strength test (Supplemental figure 2, Round 2). 1 day before and 1 day after the 14d exposure period, mice were placed into clear cylindrical plastic buckets measuring 24 cm in height and 19 cm in diameter, which were filled with 23–25°C tap water to a depth of 16-20 cm. Swim sessions were recorded over the course of a 6-minute test using a camera mounted on a tripod, which was positioned so its lens was level with the water surface in the bucket. For the pre-exposure test, an observer, unaware of the treatment conditions, analyzed the animals’ behavior. A time sampling technique was employed to categorize the predominant behavior as swimming, immobility, or climbing. Additionally, behavior was scored automatically using DBscorerV2. Manual observations by the observer were scored every 5 seconds during the final 4 minutes of the test. Meanwhile, the DBscorerV2 scoring followed best practice guidelines provided on GitHub (93). Comparisons between observer and DBscorer evaluations revealed no significant discrepancies. Consequently, DBscorerV2 was used for all subsequent swim tests.

A force transducer (Series 2 Mark-10; JLW Instruments, USA) was placed vertically and used to measure forelimb grip strength (94). Peak tensions were measured between 8AM – 11AM each testing day. Replicates were performed 3 times mouse with 1-minute breaks between. Replicates were averaged and normalized by mouse weight taken on the same morning (95). Procedurally, mice were allowed to grip the bar and the tail was slowly pulled, allowing mice to build up resistance to the force applied. In each replicate, trials resulting from a single paw grip, hind leg assistance, or a mouse body angle <30° off center relative to the tensometer were excluded and repeated.

### Brain Tissue Dissection

Under isoflurane anesthesia, all mice underwent transcardial ice-cold 0.1M PBS (pH=7.4) perfusion and subsequent steps were performed quickly. Surgical scissors were used to cut skulls from caudal to rostral along the medial edge until reaching the frontal bone anterior to bregma. Skulls were reflected rostrally to expose intact brains and olfactory bulbs. Brains were removed using a sterilized spatula and placed onto ice-cold watch glass with a sterile filter paper cover that had been soaked in ice-cold PBS (minus calcium and magnesium). Each pre-frontal cortex (PFC) was excised using a sterile razor blade and placed into a cryovial before plunging into liquid N_2_ for rapid freezing.

### Metabolomic Tissue Preparation and Analysis

Each PFC sample (~20 mg, n=6) was homogenized in an Eppendorf tube using a Bullet Blender homogenizer (Next Advance, Averill Park, NY) in 200 μL methanol:PBS (4:1, v:v, containing 1,810.5 μM 13C3-lactate and 142 μM 13C5-glutamic acid) with a final addition of 800 μL methanol:PBS (same v:v). Samples were vortexed again for 10 s, stored at −20°C for 30 min and sonicated in an ice bath for 30 min. After a centrifugation step at 14,000 RPM for 10 min (4°C), 800 μL of supernatant was transferred to a new Eppendorf tube. The samples were dried under vacuum (CentriVap Concentrator, Labconco, Fort Scott, KS). The obtained residues were reconstituted in 150 μL 40% PBS/60% acetonitrile. A quality control sample was pooled from all the study samples.

### Liquid Chromatography–Tandem Mass Spectrometry (LC–MS/MS)

Targeted LC-MS/MS techniques were similar to several recent reports (96, 97) using an Agilent 1290 UPLC-6490 QQQ-MS (Santa Clara, CA) system. Each PFC sample was injected twice: (1) 10 μL volume for negative ionization mode analysis and (2) 4 μL volume for positive ionization mode analysis. Both chromatographic separations were performed in hydrophilic interaction chromatography mode on a Waters XBridge BEH Amide column (150 x 2.1 mm, 2.5 μm particle size, Waters Corporation, Milford, MA). A flow rate of 0.3 mL/min, auto-sampler temperature of 4°C, and a column compartment temperature of 40°C were employed. The mobile phase was composed of Solvents A (10 mM ammonium acetate, 10 mM ammonium hydroxide in 95% H_2_O/5% acetonitrile) and B (10 mM ammonium acetate, 10 mM ammonium hydroxide in 95% acetonitrile/5% H_2_O). Initial 1 min isocratic elution of 90% B, decreased 40% B for 4 minutes (at t=11 min till t=15 min). The percentage of B gradually went back to 90%, to prepare for the next injection. The mass spectrometer is equipped with an electrospray ionization source. Targeted data acquisition was performed in multiple-reaction-monitoring mode. The whole LC-MS system was controlled by Agilent Masshunter Workstation software (Santa Clara, CA). The extracted MRM peaks were integrated using Agilent MassHunter Quantitative Data Analysis (Santa Clara, CA). LC-MS grade acetonitrile, methanol, ammonium acetate, and acetic acid were purchased from Fisher Scientific (Pittsburgh, PA). Ammonium hydroxide was bought from Sigma-Aldrich (Saint Louis, MO). Standard compounds were purchased from Sigma-Aldrich and Fisher Scientific. Resultant data were normalized by tissue weight before subsequent normalization steps.

### Data Analysis and Statistics

The quality control samples (inserted at 5-sample intervals during mass spectrometry) were utilized as a pooled sample group to compensate for temporal variability on the machine. Metabolomic data was analyzed using the R package MetaboAnalystR. Normality was determined via Shapiro-Wilk testing. In the event of normally distributed data, student’s t-tests were used. For non-normally distributed data, t-tests were employed on log_2_() or log_10_() transformed data. Tests were either conducted in GraphPad Prism v9.1.1, Excel (Version 2211 Build 16.0.15831.20098) or Rstudio v1.4.1564. Venn diagram was generated using an online multiple list comparison tool (98). For Venn diagrams, uncorrected student’s t-tests were performed as these data were not individually examined, but qualitatively explored as overlapping matrices. These values were input into excel, Rstudio, or downloaded directly for figure generation. Volcano plots were generated using an FDR p<0.1 and fold-change threshold >1.5. Linear mixed effects regression modeling was performed with the packages lme4, lmerTest, emmeans, multcomp, and lsmeans. Data were analyzed with fixed effects being age, exposure, and drug combination, with each metabolite being nested, and sample number called as a random effect. Results were holm-corrected before being plotted with the ggplot2, dplyr, and tidyr packages. Behavioral and other data were analyzed in GraphPad Prism using two-way ANOVAs or Student’s t-tests.

## ACKNOWLEDGEMENTS

This research was funded in part by NIH (R01AG070776 and P20GM130422).

## Supplemental Figures

**Supplemental Figure 1.**
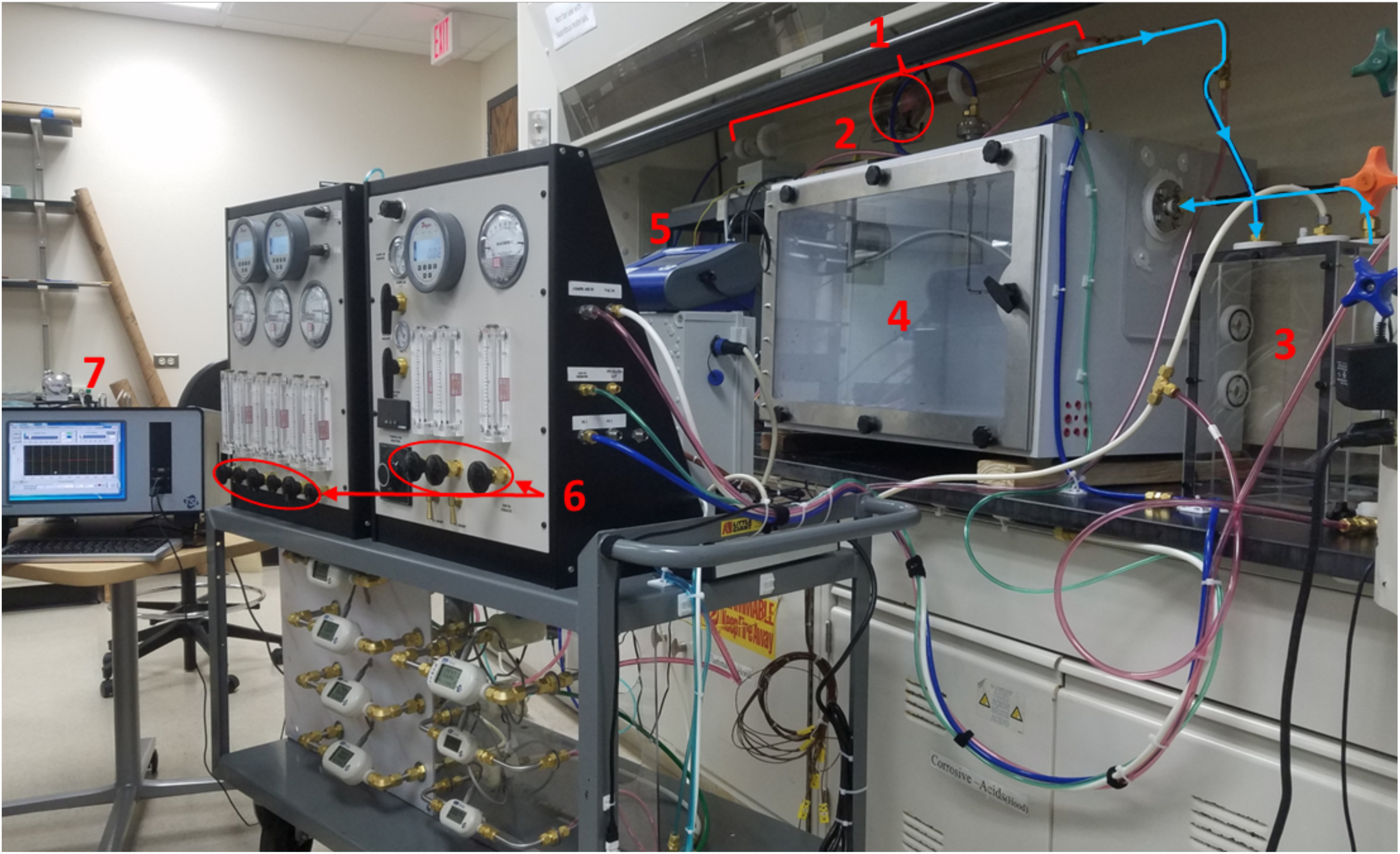
Exposure Chamber. 1) Pre-weighed combustible material is placed inside of quartz boats that are lined inside of the quartz tube. 2) Furnace that runs along the length of the quartz tube; temperature can be altered in real time or set to a desired endpoint temperature. 3) Dilution chamber where smoke can enter via suction or pressurization from the quartz tube. 4) Exposure chamber that contained mouse cages; Smoke can enter via pressurization or vacuum suction. 5) Dust Trak was used to measure particle concentration in real time. 6) Banks of dials used to alter pressure and vacuum for increased or decreased smoke exposure. 7) TSI Laser Aerosol Spectrometer was used to measure particle size distribution. wBlue arrows: direction of smoke flow.

**Supplemental Figure 2.**
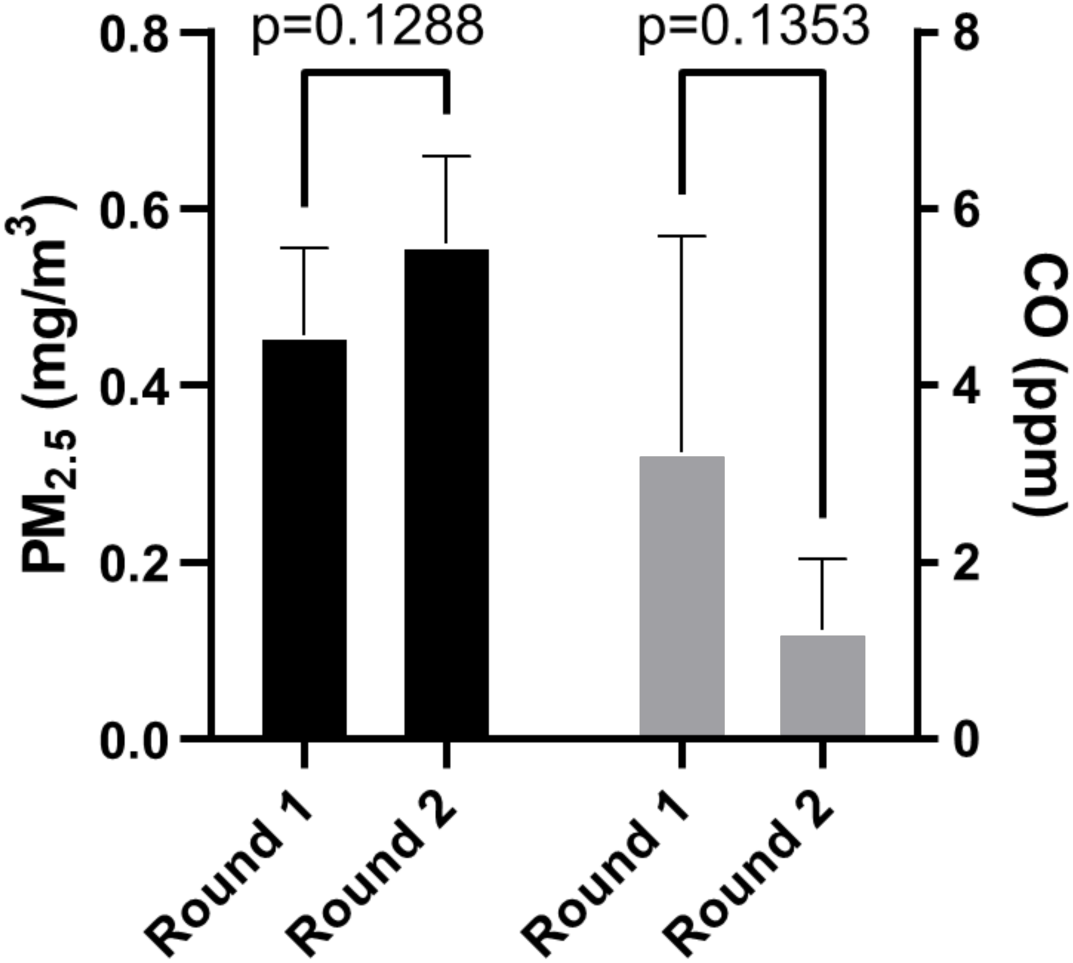
Comparison of exposures between 1^st^ cohort and 2^nd^ cohort of animals used in forced swim tests. Exposure paradigms shown as not significantly different between each other. Left axis: PM_2.5_ in mg/m^3^. Right axis: carbon monoxide in parts per million. CO levels below US EPA standards.

## Notes

**Statement of Interests:** The authors declare no conflicts of interest with the content of this manuscript.

### Competing Interest Statement

The authors have declared no competing interest.

